# New model of glucose-insulin regulation characterizes effects of physical activity and facilitates personalized treatment evaluation in children and adults with type 1 diabetes

**DOI:** 10.1101/2022.06.10.495592

**Authors:** Julia Deichmann, Sara Bachmann, Marie-Anne Burckhardt, Marc Pfister, Gabor Szinnai, Hans-Michael Kaltenbach

## Abstract

Accurate treatment adjustment to physical activity (PA) remains a challenging problem in type 1 diabetes (T1D) management. Exercise-driven effects on glucose metabolism depend strongly on duration and intensity of the activity, and are highly variable between patients. In-silico evaluation can support the development of improved treatment strategies, and can facilitate personalized treatment optimization. This requires models of the glucose-insulin system that capture relevant exercise-related processes. We developed a model of glucose-insulin regulation that describes changes in glucose metabolism for moderate- to high-intensity PA of short and prolonged duration. In particular, we incorporated the insulin-independent increase in glucose uptake and production, including glycogen depletion, and the prolonged rise in insulin sensitivity. The model further includes meal absorption and insulin kinetics, allowing simulation of everyday scenarios. The model accurately predicts glucose dynamics for varying PA scenarios in a range of independent validation data sets, and full-day simulations with PA of different timing, duration and intensity agree with clinical observations. We personalized the model on data from a multi-day free-living study of children with T1D by adjusting a small number of model parameters to each child. To assess the use of the personalized models for individual treatment evaluation, we compared subject-specific treatment options for PA management in replay simulations of the recorded data with altered meal, insulin and PA inputs.

**Author summary:** Exercise represents a cornerstone of diabetes management. Yet, many people with type 1 diabetes restrain from exercising, since it increases the risk for hypoglycemia and requires adjusted insulin treatment. The effect of exercise on blood glucose levels depends on exercise duration and intensity, but also varies strongly between individuals, making accurate adjustment a challenge. Mathematical models can help to better understand exercise physiology and to devise new treatment strategies. Here, we propose a model of glucose-insulin regulation that captures the effects of exercise on glucose metabolism and personalize it to individual children with type 1 diabetes, allowing subject-specific treatment assessment.

## Introduction

Blood glucose (BG) homeostasis maintains glucose levels within a tight range in healthy individuals, where the two main hormones involved are insulin and glucagon to lower and raise glucose levels, respectively. In type 1 diabetes (T1D), BG regulation is impeded by the autoimmune destruction of insulin-secreting *β*-cells of the pancreas [1]. The resulting lack of insulin leads to elevated glucose levels if untreated. People with T1D therefore rely on exogenous insulin either from multiple daily injections or an insulin pump together with BG monitoring to keep glucose levels stable within a target range of usually 70-180 mg/dl, with insulin requirements varying strongly between individuals. Tight glucose control is essential to avoid long-term complications such as cardiovascular disease and retinopathy from persistent hyperglycemia, or acute complications such as loss of consciousness and seizures from severe hypoglycemia.

Mathematical models of glucose-insulin regulation are a valuable tool for the in-silico evaluation of treatment strategies in T1D and play a critical role in the development of decision support and closed-loop insulin delivery systems (artificial pancreas) [2–4]. One prominent example is the UVa/Padova type 1 diabetes simulator [5] that has been approved by the FDA for preclinical testing of control algorithms for insulin treatment. While such models are typically used with hypothetical in-silico patients, a recent approach uses a personalized model to replay recorded data of individuals with T1D with altered carbohydrate (CHO) and insulin inputs, allowing subject-specific treatment assessment for improved BG control [6].

T1D treatment also needs to be adjusted to physical activity (PA), but complex PA-driven changes in glucose metabolism pose major challenges for accurate PA management. Changes occur on different time scales and strongly depend on duration and intensity of PA. Glucose demand increases drastically during the activity and insulin sensitivity remains elevated for several hours following exercise [7], leading to an increased risk for both acute and late-onset hypoglycemia. Current guidelines for treatment adjustment consider only coarse categories of glycemia, PA duration and intensity, and need further tailoring to the individual person [8, 9]. Tailoring largely relies on trial-and-error, and while PA has numerous benefits and represents a cornerstone in diabetes management [10, 11], fear of hypoglycemia restrains many people with T1D from exercising [12].

Extended models that capture exercise metabolism can help evaluate PA guidelines and treatment strategies [13] and the need for such models has long been recognized [14–16]. Roy et al. [17] proposed a PA extension of the Bergman minimal model [18], considering acute, insulin-independent effects of moderate-intensity PA on glucose uptake and production. They also included effects of liver glycogen depletion for prolonged PA. In an alternative proposal, Breton [19] studied increased glucose effectiveness and prolonged PA-driven changes in insulin sensitivity during an euglycemic hyperinsulemic clamp protocol in people with T1D. However, the effects of exercise intensity and duration on insulin action were not incorporated. Dalla Man et al. [20] integrated the model into their simulation model of the glucose-insulin system [21] in an in-silico study and added intensity- and duration-dependence, while Alkhateeb et al. [22] evaluated different variations of the Bergman minimal model and selected a model that features an increase in glucose effectiveness and insulin sensitivity. Other models have been proposed [23–26] and a virtual patient population has been generated [27] that incorporates PA [24].

However, these models have been developed under very controlled conditions, e.g. in clamp studies, have not been tailored to a T1D population, do not permit varying PA intensities or prolonged duration, or cover only a subset of the relevant processes. In addition, they often do not consider insulin and carbohydrate inputs. Hence, they are not suited for (personalized) treatment evaluation under everyday-life conditions.

Recently, Romeres et al. [28–30] and Nguyen et al. [31] conducted two elegant studies in which they evaluated exercise-induced changes in glucose utilization and endogenous glucose production, and separated and quantified insulin-dependent and –independent contributions. Incorporating their findings into models of exercise metabolism could alleviate some of the persistent problems and is useful for several reasons. A more accurate representation of exercise physiology by considering insulin-dependent and –independent effects separately facilitates prediction of exercise-driven changes in glucose levels and hypoglycemic events. In turn, this could be used to develop and evaluate improved insulin treatment strategies for PA in T1D. Furthermore, the quantification of overall glucose uptake and production rates allows to develop separate model components for each process. Previously, insulin-independent changes in glucose metabolism were often summarized in an exercise-induced increase in glucose effectiveness in PA models for T1D. As discussed by Alkhateeb et al. [22], this allows for decreasing glucose levels for moderate-intensity PA, but high-intensity PA can not be described by such models due to rising BG levels. In addition, it is difficult to incorporate liver glycogen depletion that affects the rate of glucose production for prolonged PA.

Here, we utilize these newly available data and develop a glucose-insulin regulation model for exercise that explicitly considers insulin-dependent and -independent effects on glucose uptake and production, and allows realistic full-day simulations and personalized replay simulations. The model captures the acute and prolonged changes in glucose metabolism during PA and subsequent recovery for moderate-to high-intensity exercise, and considers CHO intake and insulin injections. We first calibrate the model for a healthy population, before adjusting relevant parameters to people with T1D. We validate the model on independent data from increasingly complex scenarios including PA, insulin and CHO intake.

We show how exercise duration, intensity and time of day alter BG dynamics in full-day simulations. As a main result, we demonstrate that our model can describe real-world data of individual patients. We personalize the model on data from children with T1D recorded in a free-living observational study, using only data from sensors readily available during everyday-life. We then perform replay simulations of the original scenarios with altered meal, insulin and PA inputs to evaluate different treatment strategies and PA effects on the individual subject level.

## Methods

### Development of a Glucoregulatory Model Including Physical Activity

Our proposed model is outlined in Fig 1 and comprises a simple core model extended by meal intake and insulin injections. Accelerometer counts *AC* quantify the input for PA processes affecting glucose regulation.

**Fig 1.**
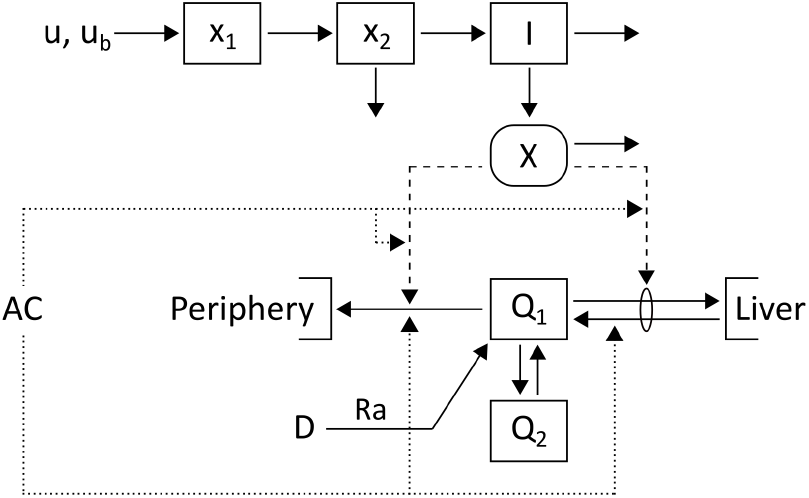
Schematic of the glucose-insulin model. Glucose, *Q*_1_, and insulin, *I*, dynamics are described using a two-compartment model [32]. Extensions capture plasma insulin kinetics after subcutaneous injection *u* with basal insulin infusion rate *u*_*b*_ [33] and glucose appearance with rate *Ra* after a meal *D* [34]. PA is measured via accelerometer counts *AC* and leads to changes in glucose metabolism indicated by dotted lines.

We use the Cobelli two-compartment minimal model [32] to describe glucose-insulin regulation at rest and extend it to incorporate PA-driven changes in glucose metabolism:

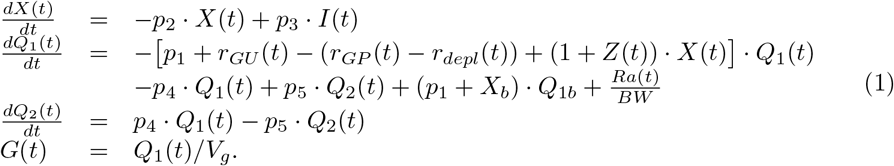

The two glucose compartments *Q*_1_ and *Q*_2_ [mg/kg] represent glucose mass in plasma and in a remote compartment, respectively. Plasma insulin *I* [*μ*U/ml] promotes the disappearance of plasma glucose into liver and tissue, and suppresses hepatic glucose production via the dynamic state *X* [1/min]. The constants *Q*_1*b*_ [mg/kg] and *X*_*b*_ = *p*_3_*/p*_2_ · *I*_*b*_ [1/min] provide the basal levels of plasma glucose and state *X*, respectively, with the basal plasma insulin level *I*_*b*_ [*μ*U/ml]. The term *p*_3_*/p*_2_ represents insulin sensitivity and *p*_1_ describes glucose effectiveness. The rate parameters *p*_4_ and *p*_5_ quantify the exchange between the two glucose compartments. Glucose appearance from meals is described by *Ra* [mg/min] and scaled with bodyweight, *BW* [kg]. The rates *r*_*GU*_ [1/min] and (*r*_*GP*_ − *r*_*depl*_) [1/min] provide the exercise-induced insulin-independent increase in glucose uptake (GU) and production (GP), respectively, while (1 + *Z*) captures a PA-driven rise in insulin sensitivity. Finally, *G* [mg/dl] is the plasma glucose concentration and *V*_*g*_ [dl/kg] the glucose distribution volume.

#### Measure of exercise intensity and duration

We consider accelerometer (AC) counts to capture movement and link them to PA intensity *Y* [counts/min] following previous approaches [17, 19, 23]:

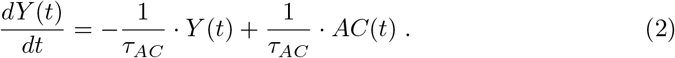

The delay *τ*_*AC*_ [min] allows initial adaptation to PA.

We also track PA duration *t*_*P A*_ [min], integrated AC count *PA*_*int*_ [counts] and time spent at high intensity *t*_*h*_ [min]:

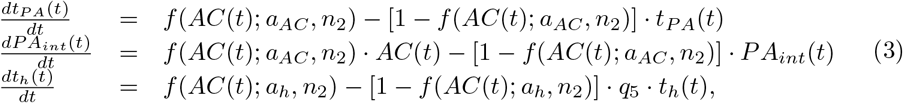

where transfer functions *f* (*AC*; *a*_*AC*_, *n*_2_) and *f* (*AC*; *a*_*h*_, *n*_2_) capture the transition in AC count from rest to exercise, respectively from moderate to high intensity, with

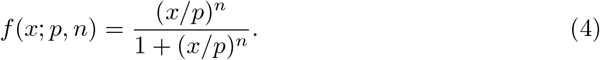

The use of transfer functions to introduce exercise-related changes was previously proposed by Breton [19].

#### Insulin sensitivity

Insulin sensitivity increases during exercise and stays elevated afterwards for up to 48 hours to replete liver glycogen stores [35]. Previous studies have further established that the increase depends linearly on PA intensity and duration [20, 22], and we consequently describe this rise *Z* by

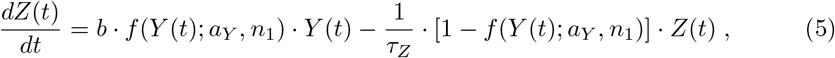

where *f* (*Y* ; *a*_*Y*_, *n*_1_) defines the minimal intensity *Y* considered as PA, parameter *b* [1/count] specifies the proportional rise, and *τ*_*Z*_ [min] the time for insulin sensitivity to return to its baseline level.

#### Insulin-independent glucose uptake and production

Glucose demand by active muscles increases acutely during PA and glucose uptake from plasma is upregulated. Simultaneously, hepatic glucose production by gluconeogenesis and glycogenolysis increases to maintain plasma glucose levels [36]. These processes are linear in PA intensity [17]. We therefore define the insulin-independent rise in GU (*r*_*GU*_ [1/min]) and GP (*r*_*GP*_ [1/min]) rates as

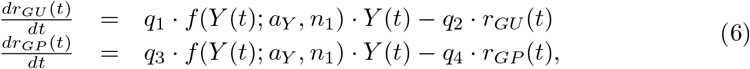

where *q*_*i*_ are rate parameters.

#### Glycogen depletion

Liver glycogen stores may deplete during prolonged PA and GP cannot be maintained by gluconeogenesis alone, causing a drop in glucose levels [37]. We follow Roy et al. [17] and assume that glycogen stores deplete in proportion to exercise intensity and duration. The time *t*_*depl*_ [min] to depletion determined from the integrated AC count and PA duration is then given by:

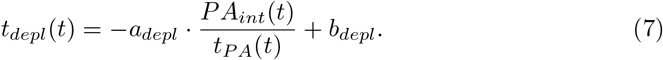

After depletion sets in, we allow a drop in GP rate, *r*_*depl*_ [1/min], defined by

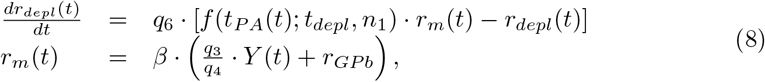

where the transfer function *f* (*t*_*PA*_; *t*_*depl*_, *n*_1_) indicates whether exercise time exceeds *t*_*depl*_ and *q*_6_ is a rate parameter. The maximum decrease *r*_*m*_ [1/min] in GP is the sum of the basal resting GP rate, *r*_*GP b*_, and the PA-driven GP rate at steady state, *q*_3_*/q*_4_ · *Y* (*t*), scaled by the proportion of net hepatic glucose production attributed to glycogenolysis, *β*.

#### High-intensity exercise

During high-intensity 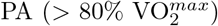, GP may (initially) exceed GU and result in rising plasma glucose levels due to an increase in catecholamines and cortisol [38]. We mimic the drastic rise in GP by modulating parameters *q*_3_ and *q*_4_ between low- (subscript *l*) and high-intensity (subscript *h*) values

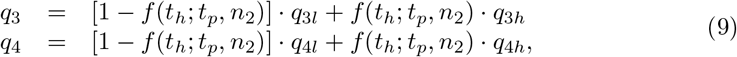

where we use the transfer function *f* (*t*_*h*_; *t*_*p*_, *n*_2_) to smoothly transition between the two exercise regimes.

### Model Extensions for Full-Day Simulations

To enable full-day simulations, we further include existing models to provide plasma insulin concentration after insulin injections and rate of glucose appearance after meals, and use these as inputs to the exercise model.

#### Insulin kinetics

We use a model with two subcutaneous compartments of insulin masses *x*_1_ and *x*_2_ [*μ*U] and a plasma insulin compartment *I* [*μ*U/ml] to model plasma insulin after a subcutaneous injection [33]:

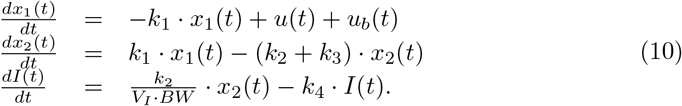

Insulin is injected into *x*_1_, with *u* [*μ*U/min] and *u*_*b*_ [*μ*U/min] defining the rates of correction and basal insulin infusion, respectively. *V*_*I*_ [ml/kg] is the insulin distribution volume and *k*_*i*_ are rate parameters. We estimated the model parameters from insulin measurements obtained after a subcutaneous injection of 0.3 U/kg insulin aspart [39] (Sec B in S1 File).

#### Carbohydrate absorption

We describe the glucose appearance rate *Ra* [mg/min] after a meal with carbohydrate content *D* [mg] with an established model [34]:

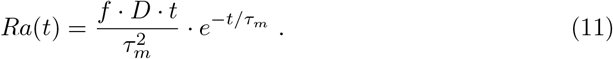

A fraction *f* of glucose is absorbed into plasma and the time constant *τ*_*m*_ [min] characterizes the time-of-maximum appearance rate. We determine *f* and *τ*_*m*_ individually for each meal (see below).

### Model Calibration

We obtained parameter values from literature or physiological knowledge when feasible and estimated the remaining parameters from published data. We followed a stepwise approach for parameter estimation and calibrated a population-average model on data of healthy subjects acquired during exercise, before adjusting parameters to describe glucose metabolism and PA effects in people with T1D (Table 1).

**Table 1.** Model parameters. Healthy: healthy subjects. V1–V3: T1D subjects under euglycemia low-insulin (V1), euglycemia high-insulin (V2), and hyperglycemia low-insulin (V3) clamp conditions [30, 40].

| Parameter | Healthy | T1D |  |  | Unit |
| --- | --- | --- | --- | --- | --- |
|  |  | V1 | V2 | V3 |  |
| Glucose-Insulin Regulation |  |  |  |  |  |
| $p_1$ | 0.017 | 0.0126 | 0.0214 | 0.0114 | 1/min |
| $p_2$ | 0.0351 | 0.0228 | 0.0228 | 0.0228 | 1/min |
| $p_3$ | $2.47 \cdot 10^{-5}$ | $2.78 \cdot 10^{-5}$ | $2.98 \cdot 10^{-5}$ | $2.19 \cdot 10^{-5}$ | ml/(μU·min<sup>2</sup>) |
| $p_4$ | 0.058 | 0.058 | 0.058 | 0.058 | 1/min |
| $p_5$ | 0.0885 | 0.0885 | 0.0885 | 0.0885 | 1/min |
| $V_g$ | 1.289 | 1.289 | 1.289 | 1.289 | dl/kg |
| Insulin Sensitivity |  |  |  |  |  |
| $\tau_{AC}$ | 5 | 5 | 5 | 5 | min |
| $\tau_Z$ | 600 | 600 | 600 | 600 | min |
| $b$ | $1.68 \cdot 10^{-6}$ | $3.64 \cdot 10^{-6}$ | $1.59 \cdot 10^{-6}$ | $1.55 \cdot 10^{-6}$ | 1/count |
| Glucose Uptake and Production |  |  |  |  |  |
| $\alpha$ | 0.27 | 0.27 | 0.27 | 0.27 | dimensionless |
| $q_1$ | $1.93 \cdot 10^{-7}$ | $6.46 \cdot 10^{-7}$ | $2.88 \cdot 10^{-6}$ | $5.23 \cdot 10^{-7}$ | 1/(count·min) |
| $q_2$ | 0.066 | 0.0617 | 0.2682 | 0.0778 | 1/min |
| $q_{3l}$ | $1.15 \cdot 10^{-7}$ | $4.46 \cdot 10^{-7}$ | $1.33 \cdot 10^{-7}$ | $1.60 \cdot 10^{-7}$ | 1/(count·min) |
| $q_{4l}$ | 0.051 | 0.0705 | 0.0299 | 0.0669 | 1/min |
| $q_{3h}$ | - | - | - | $5.87 \cdot 10^{-7}$ | 1/(count·min) |
| $q_{4h}$ | - | - | - | 0.056 | 1/min |
| $q_5$ | - | - | - | 0.03 | 1/min |
| Glycogen Depletion |  |  |  |  |  |
| $\beta$ | 0.76 | - | - | - | dimensionless |
| $q_6$ | 0.1 | - | - | - | 1/min |
| $a_{depl}$ | 0.0108 | - | - | - | min<sup>2</sup>/count |
| $b_{depl}$ | 180.6 | - | - | - | min |
| Transfer Functions |  |  |  |  |  |
| $a_Y$ | 1500 | 1500 | 1500 | 1500 | counts/min |
| $a_{AC}$ | 1000 | 1000 | 1000 | 1000 | counts/min |
| $a_h$ | 5600 | 5600 | 5600 | 5600 | counts/min |
| $t_p$ | 2 | 2 | 2 | 2 | min |
| $n_1$ | 20 | 20 | 20 | 20 | dimensionless |
| $n_2$ | 100 | 100 | 100 | 100 | dimensionless |

#### Parameter determination for healthy subjects

We used the original parameter values of the two-compartment minimal model [32] and explicitly included the effect of basal insulin on glucose. To calibrate the exercise model, we separated parameters into process-specific sets and individually estimated these on data sets acquired during the corresponding exercise modes using least squares regression.

We set the delay parameter *τ*_*AC*_ to 5 min [19, 20] and chose a time constant *τ*_*Z*_ of 600 min [20] such that insulin sensitivity stays elevated for up to 48 hours in accordance with literature reports [35].

We obtained the increase in insulin sensitivity during PA (parameter *b*) from measurements of the insulin-dependent rate of glucose disappearance during rest and 100 min of cycling at 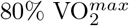 [41]. We converted 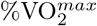 to accelerometer count using

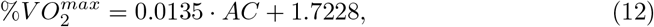

estimated from simultaneous AC count and 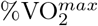 measurements for different types and intensities of PA [42].

We estimated the insulin-independent GU and GP parameters *q*_1_, *q*_2_, *q*_3*l*_ and *q*_4*l*_ from total GU and GP rates measured during 60 min of PA at 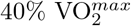 [43] (Fig S1a in S1 File). We distinguished between resting and exercise-driven contributions by separating the net rate of glucose change into endogenous glucose production and glucose uptake:

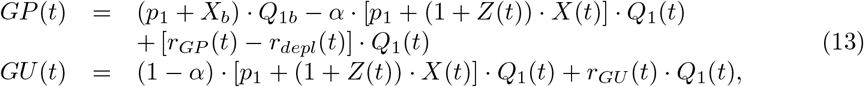

where we determined *α* from measurements at rest (*Z* = *r*_*GU*_ = *r*_*GP*_ = *r*_*depl*_ = 0), and assumed that the prolonged exercise-driven change in insulin sensitivity affects both GP and GU as found in Romeres et al. [29, 30].

We determined the time until hepatic glycogenolysis decreases due to glycogen depletion from reported depletion times for different intensities [37] (parameters *a*_*depl*_ and *b*_*depl*_). We estimated glycogen depletion parameters *β* and *q*_6_ from plasma glucose measurements [44] recorded during 180 min of cycling at 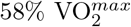, where we restricted *q*_6_ to 0.1 min^−1^ to avoid an overshoot in GP after PA and kept the remaining parameters fixed (Fig S1b in S1 File).

Finally, moderate-intensity PA is defined by AC counts above 2296 counts/min [42] and we enforced the transition from rest to PA between 1000 and 2000 counts/min with parameters *a*_*Y*_ = 1500 counts/min and *n*_1_ = 20. Accordingly, we defined *a*_*AC*_ = 1000 counts/min and *n*_2_ = 100 to track duration and AC count immediately from the start of PA. High-intensity PA commences at 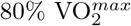 (5800 counts/min) [38], and we set *a*_*h*_ = 5600 counts/min and *t*_*p*_ = 2 min for a transition between intensity regimes at 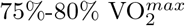.

#### Adjustment of model parameters to T1D

To re-calibrate the exercise model to persons with T1D, we relied on the study by Romeres et al. [30, 40], where people with T1D performed 60 min of exercise at 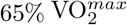 during a glucose clamp under three different glucose and insulin conditions (V1: euglycemia – low insulin, V2: euglycemia – high insulin, V3: hyperglycemia – low insulin). Plasma glucose and insulin concentrations were measured and glucose disappearance and production rates were determined from recorded data. We estimated parameters defining insulin-independent (*p*_1_, *q*_1_-*q*_4*l*_) and -dependent (*p*_3_, *b*) contributions to GU and GP at rest and during PA. For further details on the estimation procedure, see Section A.2.1 in S1 File. Additionally, we computed confidence intervals for all parameter estimates from profile likelihoods to determine practical identifiability (Sec A.2.2 in S1 File).

We estimated the high-intensity exercise parameters *q*_3*h*_ and *q*_4*h*_ from interstitial glucose measurements [45] of people with T1D performing 45 min of 4 min intervals at 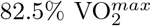 using least squares regression (Fig S4 in S1 File). We introduced the parameter *q*_5_ = 0.03 min^−1^ to prevent a switch to low-intensity parameters during recovery. For the remaining parameters, we used the values determined for the hyperglycemia - low insulin condition (V3) of the previously discussed data, as the high-intensity activity was recorded under comparable conditions.

### Model Validation

We used independent data from six additional studies covering a range of exercise intensities and durations for validating our model. Importantly, several studies include pre-exercise meal intake and insulin bolus injections as well as different insulin reduction strategies. This allowed us to validate the individual model parts and their interplay in the full model. The data sets are the following:

1. In a study by Rabasa-Lhoret et al. [46], participants with T1D performed exercise at three intensities (25%, 50% and 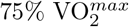) for different durations (30 and 60 min). Breakfast with 75g of CHO and varying insulin bolus sizes (25%, 50% and 100% of typical dose) was consumed 90 min prior to PA. Plasma glucose was measured.
2. In a second study conducted by Maran et al. [47], participants with T1D performed 30 min of exercise at 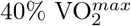. The changes in plasma glucose and insulin concentrations were recorded.
3. Participants with T1D exercised for 45 min at 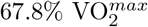 in a study presented by Iscoe and Riddell [48], where the change in interstitial glucose levels was measured.
4. The effect of basal insulin suspension during exercise was studied by Zaharieva et al. [49]. Exercise was performed for 40 min at 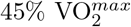 and the change in plasma glucose concentration was assessed.
5. In a study by Dubé et al. [50], exercise was performed for 60 min at 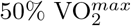, 2h after lunch including a pre-meal insulin bolus. Plasma glucose levels were monitored and participants did or did not consume a drink containing 30g of glucose 15 min pre-exercise.
6. Healthy subjects cycled for 240 min at 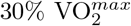 in a study conducted by Ahlborg et al. [51] and plasma glucose was measured. For this PA duration, glycogen depletion affects GP and subsequently glucose levels.

For studies (1)–(5), we used the parameter values obtained for the T1D population and allowed only slight adjustments to accommodate differences in study population and experimental conditions. For study (6), we used the parameter values of the healthy population and kept them unchanged. The resulting glucose trajectories are shown in Fig 2 and Fig S6 in S1 File. We observe good agreement between data and model predictions across all studies.

**Fig 2.**
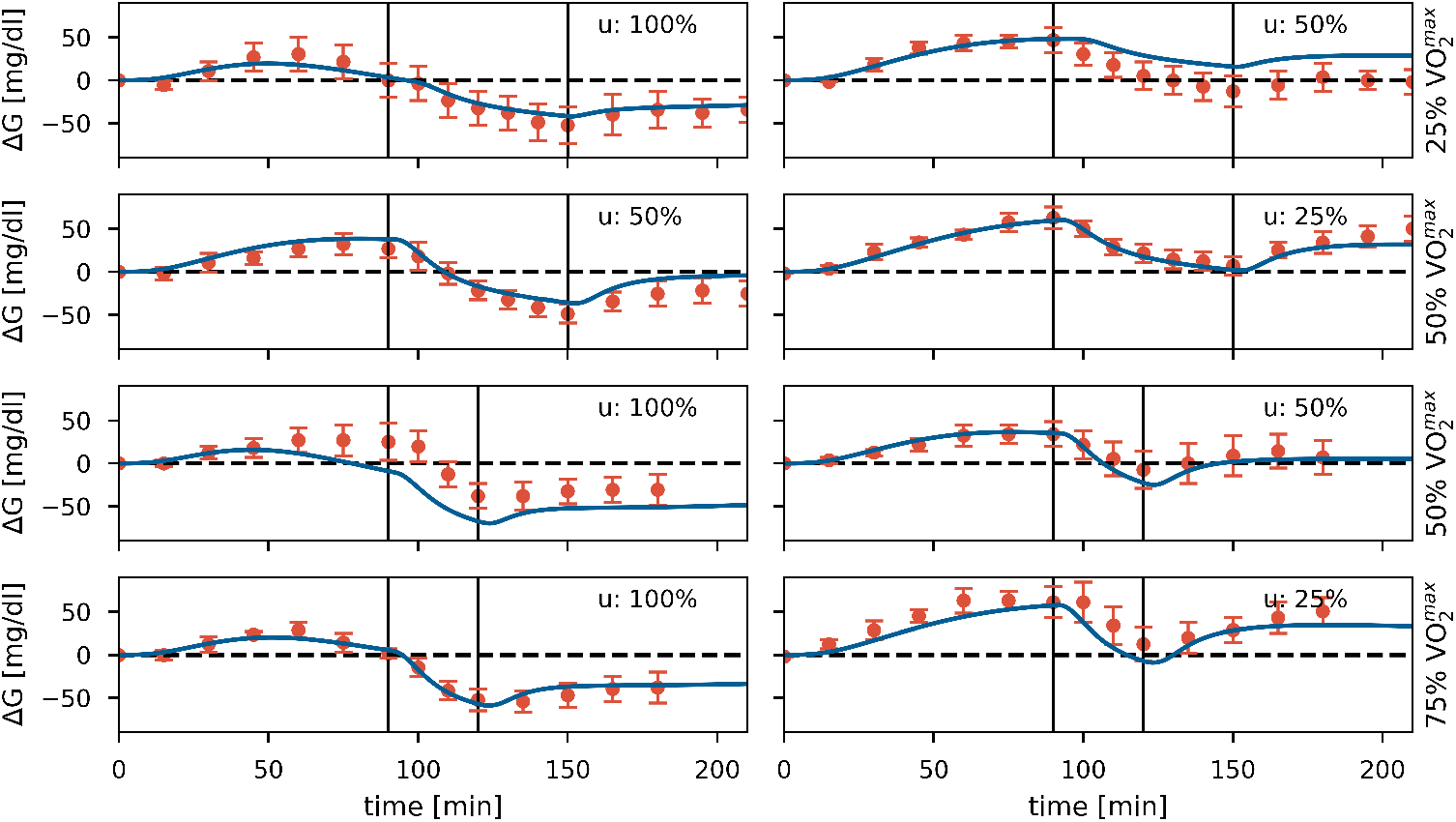
Model validation. Data [46] and model predictions for validation study (1). PA sessions are marked by vertical lines. PA was performed at different intensities (25%, 50% and 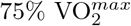) for durations of 30 or 60 min. A meal was consumed 90 min prior to PA, with a meal insulin bolus *u* of 100%, 50% or 25% of the full dose. Parameters *p*_1_ = 0.0197, *p*_3_ = 2.59 10^−5^, *f* = 2 and *τ*_*m*_ = 109 min were determined from glucose levels at rest and we applied T1D exercise parameters (V2).

## Results

### Effect of Physical Activity in Full-Day Simulations

We evaluate our model’s performance in full-day simulations for a range of PA scenarios to confirm that it reproduces clinical knowledge. We define a standard day consisting of three meals and corresponding insulin bolus injections. We include a PA session in the morning or afternoon and consider different intensities (30%, 60% and 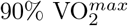) and durations (30, 60 and 180 min) (Fig 3).

**Fig 3.**
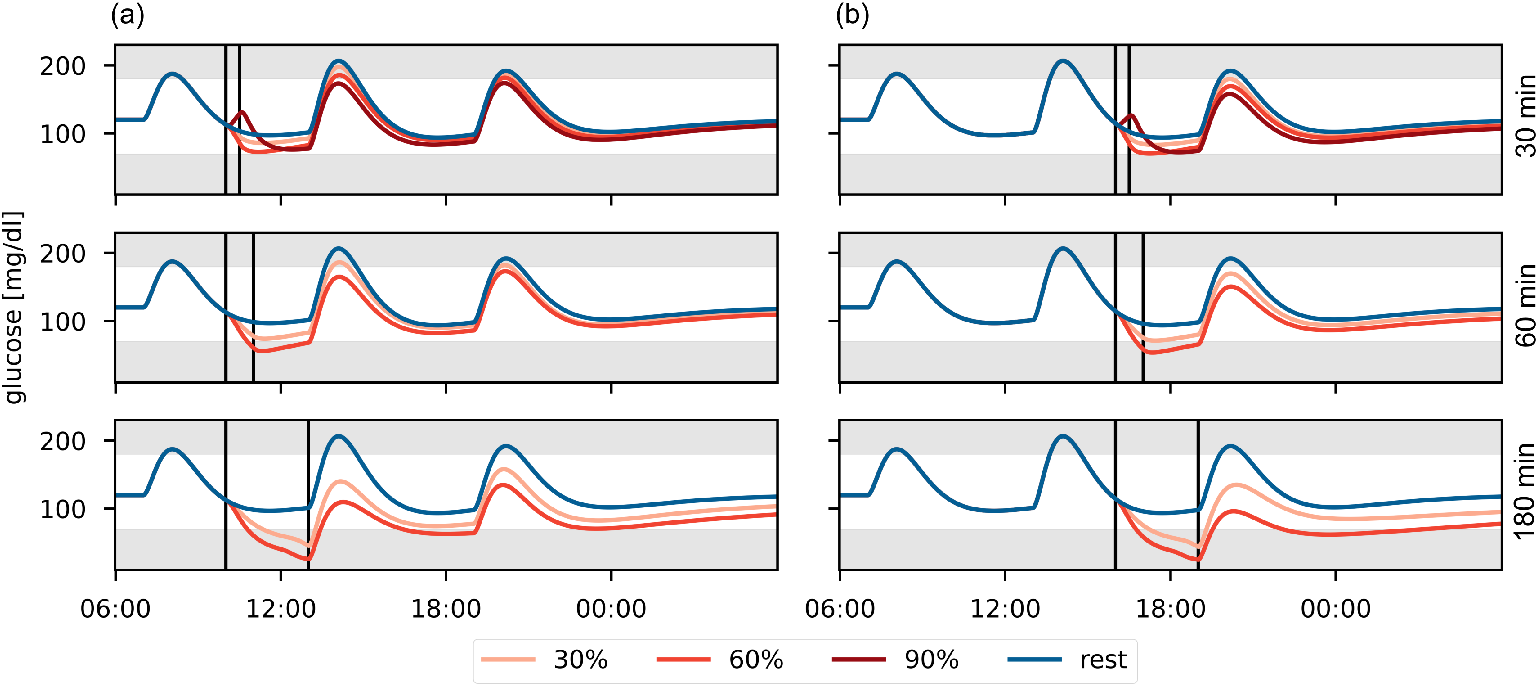
Comparison of glucose trajectories for different PA types in a full-day simulation. PA is performed at 30%, 60% and 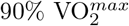 for 30, 60 and 180 minutes (a) in the morning, or (b) in the afternoon. The PA session is marked by vertical lines. Meals are eaten at 7:00, 13:00 and 19:00 containing 40g, 60g and 50g CHO, respectively.

During moderate-intensity PA, BG levels decrease with increasing intensity and duration [8] and drop even further once glycogen stores deplete [7]. In contrast, BG levels may rise [8] during high-intensity PA, which can provide protection against acute hypoglycemia [52], but the risk for late-onset hypoglycemia still increases with higher PA intensity and duration [47, 53]. Time of day also affects the risk for nocturnal hypoglycemia, which is higher for afternoon-compared to morning-PA [54].

Our model accurately reflects the duration- and intensity-dependence for moderate- and high-intensity PA. As expected, BG levels increase during high-intensity PA, but drop below those of moderate-intensity PA after the activity. We also find lower nocturnal BG levels following afternoon-compared to morning-PA. Our model thus reproduces clinical observations regarding hypoglycemia risk for a range of different PA scenarios.

### Model Personalization on Data from Children with T1D

To establish our model’s capability to describe individual subject data, we personalize the model on multi-day at-home data from five children aged 8–14 with T1D [55] (‘DiaActive’ study, ethics approval no. 341/12, Ethics Commission Cantons of Basel, February 14, 2013; written formal consent was obtained from the parent/guardian for each study participant). For each participant, interstitial glucose levels were measured by continuous glucose monitoring (CGM), exercise was monitored using an accelerometer, and CHO content and timing of meals as well as timing and dosing of insulin injections were self-reported in logbooks. We provide participant characteristics in Section E.2.1 and discuss data preparation in Section E.2.2 in S1 File.

To personalize the model to each participant, we followed a strategy presented by [6] and determined subject-specific parameter values for insulin sensitivity (*p*_3_) and meal parameters (*f* and *τ*_*m*_) using least squares regression. We also estimated glucose effectiveness (*p*_1_) and the basal glucose concentration *G*_*b*_. We computed the basal insulin level *I*_*b*_ based on the basal insulin infusion rate *u*_*b*_, and kept the remaining parameter values, including all exercise-related parameters, at their previously determined population-average level (Table 1, V1). We confirmed local structural identifiability of the re-calibrated parameters using the STRIKE-GOLDD toolbox [56]. Furthermore, we established our model’s capacity for personalization and replay simulations with altered meal and insulin inputs–but no PA–using the UVa/Padova simulator (Python implementation [57]) (Sec E.1 in S1 File). Note that we forewent the deconvolution step originally proposed to address further model mismatch.

We consider two–not necessarily consecutive–24 hour periods for each of the five children. We estimate parameters *p*_1_ and *p*_3_ on data from the first day and keep these values for the second day to confirm that the personalized models generalize to new scenarios. We estimate basal glucose for each day, and estimate meal parameters independently for each meal to account for inaccuracies in the self-reported meal sizes and for different meal compositions.

The individual model fits show very good overall agreement with the recorded data and we only observe a small number of non-explainable glucose excursions, which we attribute to unrecorded meals or unknown residual dynamics carried over from the previous day (Fig 4, and Fig S8 and Tables S3-S5 in S1 File). We quantified the model fits using the root mean square difference (RMSD) and the mean absolute relative difference (MARD) (Table 2). We reach commonly used targets of RMSD below 25 mg/dl [58] and MARD below 10% [6] in most cases.

**Fig 4.**
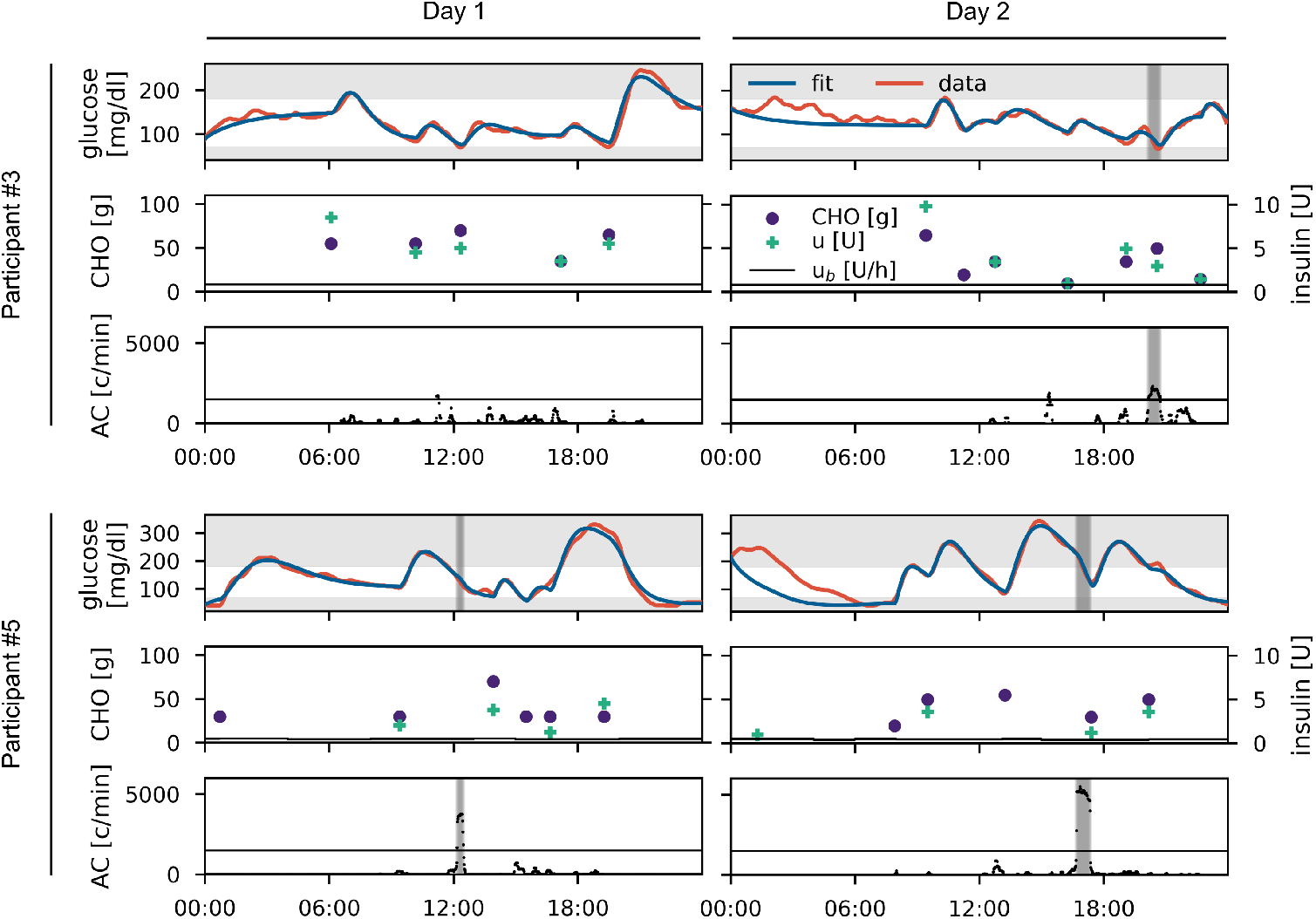
Data and personalized model for study participants #3 and #5 for two days each. For each day, recorded (red) and fitted (blue) glucose data are shown in the upper panel. Carbohydrate (purple) and insulin (green) inputs are shown in the middle panel. Accelerometer counts are shown in the lower panel with periods of physical activity highlighted in grey. The remaining participants are shown in Fig S8 in S1 File.

**Table 2.** Evaluation of personalized model fits. Unexplained glucose excursions are excluded and results for the full 24h period are given in brackets in these cases.

| Participant | # 1 |  | # 2 |  | # 3 |  | # 4 |  | # 5 |  |
| --- | --- | --- | --- | --- | --- | --- | --- | --- | --- | --- |
|  | day 1 | day 2 | day 1 | day 2 | day 1 | day 2 | day 1 | day 2 | day 1 | day 2 |
| RMSD [mg/dl] | 11.4 | 12.1 | 3.7 | 6.6 | 9.3 | 15.8 | 11.0 | 16.8 | 12.8 | 9.0 |
|  |  | (84.6) | (34.2) | (36.6) |  |  |  |  |  | (43.1) |
| MARD [%] | 6.7 | 10.3 | 2.9 | 5.4 | 5.9 | 7.6 | 5.3 | 10.7 | 10.1 | 4.9 |
|  |  | (22.0) | (7.8) | (15.7) |  |  |  |  |  | (15.9) |

### Replay Simulations using Personalized Models

Next, we use the personalized models to demonstrate their potential in replay simulations, a promising approach for comparing and evaluating subject-specific treatment strategies in-silico, on two 24 hour episodes selected from our data.

We first consider day 1 of participant #3 with no PA and replay these data with changes in lunch size or with an altered meal bolus (Fig 5a). As expected, a larger meal or lower bolus increase BG levels, while a smaller meal or larger bolus lead to a corresponding reduction. Differences between these treatments reduce over time and virtually vanish after dinner.

**Fig 5.**
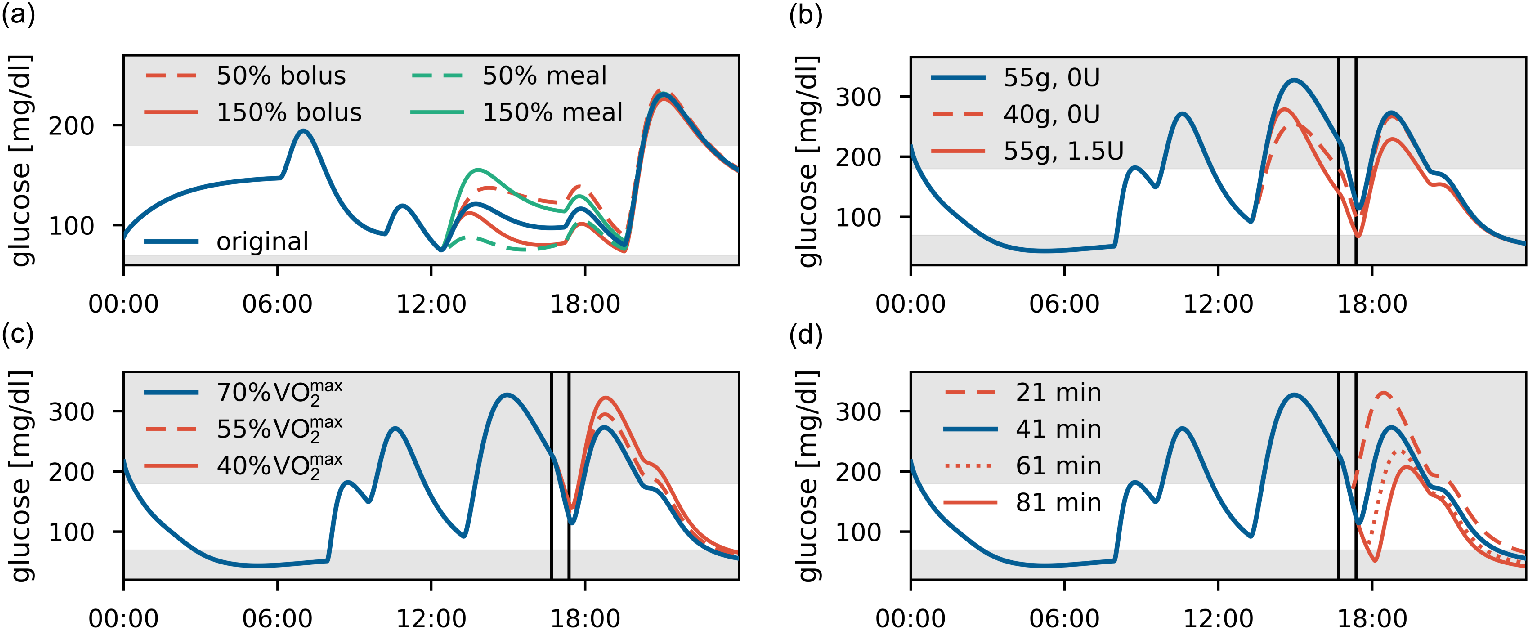
Replay simulations. (a) Participant #3, day 1. Variations in meal size and insulin dose at lunch to 50% and 150% of their original size. (b)-(d) Participant #5, day 2, with a PA session marked by vertical lines. The original glucose trajectory is shown in blue. (b) Meal or bolus adjustment for pre-PA meal. (c) Alterations in PA intensity from 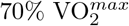 to 55% and 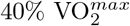. (d) Alterations in PA duration from 41 min to 21, 61 and 81 min, with the post-exercise meal following directly after the session.

For our second scenario, we use the data for day 2 of participant #5, who exercises for 41 min at almost constant intensity of 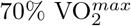 in the afternoon. Likely in anticipation of the planned PA session, the participant used no insulin bolus for the preceding meal and commenced the session in hyperglycemia. We therefore ask if an alternative treatment decision might have led to a more favorable BG trajectory, and consider reducing the pre-PA meal size from 55g to 40g CHO or to administer an insulin bolus of 1.5U (Fig 5b). The replay simulation indicates that a smaller pre-PA meal could have been favorable, reducing hyperglycemia before PA without increasing the risk of hypoglycemia during the activity.

We also use this scenario to study the effect of exercise intensity and duration. First, we replay the scenario with lower PA intensities (Fig 5c). Notably, BG trajectories only start to diverge substantially after the post-PA meal as insulin sensitivity remains elevated for several hours during recovery, and further measures to avoid post-PA hyperglycemia might be required. Next, we consider varying PA duration (Fig 5d). The effect of elevated insulin sensitivity is clearly visible as the BG trajectories stay separated for the remaining simulation time, and results suggest that no additional adjustments are necessary to protect against hypoglycemia for exercise up to an hour.

## Discussion

Mathematical models are a valuable tool to develop and evaluate treatment strategies for T1D in-silico. Finding accurate individual treatment adjustments for physical activity remains a complex process that could be facilitated by in-silico treatment evaluation, but comprehensive models including all relevant aspects of exercise metabolism suitable for this task are currently lacking.

PA-driven changes in glucose metabolism act on different time scales and require different treatment adjustments. In particular, insulin-independent processes affect BG levels mainly during PA, while insulin-dependent effects are the main cause for late-onset hypoglycemia and need to be considered for several hours post-PA. BG levels often fall during moderate-intensity PA when GU exceeds GP, while a drastic rise in GP during high-intensity PA can in contrast cause rising BG levels. It is therefore crucial to incorporate all relevant exercise processes in a PA model to study PA management in-silico.

In this work, we presented a model of glucose-insulin regulation in T1D that covers acute insulin-independent changes in GU and GP during PA, and the prolonged PA-induced rise in insulin sensitivity. We considered PA of moderate to high intensities, and accounted for depletion effects during prolonged exercise. We suggested the use of transfer functions to switch between these different exercise regimes, with the aim to keep the model compact without affecting the individual PA processes. The model includes modules for insulin bolus injections and meal intake as additional inputs to capture all aspects of daily life and diabetes management, allowing simulation of realistic scenarios.

We proposed a stepwise approach for model calibration, estimating parameters of the different model components separately on corresponding population-average data from healthy subjects. While the full model is not identifiable—a common problem for models of the glucose-insulin system—this allowed us to quantify individual contributions of the different PA-related processes accurately. Next, we adjusted the full model to a T1D population and computed profile likelihoods to determine practical parameter identifiability. We validated the model on independent data sets covering PA of different intensities and duration, and PA in conjunction with CHO intake and insulin injections. The model accurately predicts the resulting BG trajectories in all cases, demonstrating the feasibility of stepwise model identification and its potential for calibrating complex T1D models. Additionally, we evaluated the model’s prediction capabilities in full-day simulations with a range of PA scenarios against clinical knowledge.

The presented model structure is consistent with literature reports studying exercise-induced changes in glucose metabolism. Studies demonstrated that glucose utilization increases with exercise intensity for healthy subjects [59, 60] and for people with T1D [61]. It was shown by [30] and [31] that this increase can be separated into insulin-dependent and –independent contributions. In particular, they confirmed that insulin-mediated GU increases gradually during the activity and remains elevated for several hours post-PA, while non-insulin-mediated GU increases rapidly at PA onset and drops to its baseline level immediately after. [31] did not find intensity-dependence for GU, but discuss that this might have been caused by PA intensities that were not sufficiently different or by varying levels of fitness between participants.

Similarly, it was shown that endogenous glucose production increases with exercise intensity to counteract the rise in GU [31, 59–62]. In contrast to its effect on GU, insulin suppresses GP. [28] found that the PA-driven rise in GP is consequently inhibited in people with T1D with hyperinsulinemia, and identified a delayed effect of insulin on GP [29]. In contrast, [31] did not find insulin-mediated changes in GP. In this work, we followed the findings of [29], since GU and GP rates were estimated in a model-independent way, and our model is able to describe their data well when including a PA-driven, elevated effect of insulin on GP.

Furthermore, the rate of hepatic glycogenolysis during PA increases linearly with intensity [37], supporting our assumption that the time until depletion occurs decreases in proportion to PA intensity. However, we estimated depletion parameters of the model only on data from healthy subjects, and data from individuals with T1D are required to validate the model for this population. [62] observed that differences in GP between healthy and T1D subjects arise from varying contributions of gluconeogenesis, and that glycogenolysis at rest and for different PA intensities is similar for both populations. Hence, we believe that our model assumptions also hold for individuals with T1D and that predictions are therefore qualitatively correct for this population.

During high-intensity PA, counterregulatory hormones such as catecholamines and cortisol are upregulated. They are associated with increased hepatic GP that exceeds GU, and thus lead to (initially) rising BG levels during exercise [9, 38]. The rise in BG levels persists only while these hormone levels are elevated, and is followed by several hours with an increased risk for hypoglycemia [63]. We incorporated the drastic rise in GP in our model and were able to accurately reflect the resulting glucose dynamics. However, we were unable to perform an independent validation for this scenario due to lack of additional data.

Here, we only considered aerobic exercise of moderate to high intensity. We did not incorporate anaerobic exercise that is encountered for example in high-intensity interval or strength training. Anaerobic exercise can cause different trends in glucose levels for people with T1D [15], and it would be useful to integrate this modality into a PA model.

We applied our model to evaluate subject-specific treatment strategies in-silico based on model personalization and replay simulations. First, we validated this approach for altered meal and insulin inputs–but without exercise–against the UVa/Padova simulator. We then personalized our model to several children with T1D by adjusting a small number of parameters to each child, and accurately reproduced their glucose data recorded under real-life conditions. We presented examples of replay simulations from these personalized models to study subject-specific treatment alternatives and PA effects. Our results provide a promising proof-of-principle for adjusting treatment strategies to the individual person to improve PA management. The approach only requires data easily available in everyday settings from CGM devices and activity trackers, and we therefore expect that it also applies to the challenging case of unplanned and unstructured PA typical for children.

We anticipate that our model could be used in practice to describe and simulate blood glucose levels and to predict hypoglycemia associated with PA. Model personalization allows replay of recorded data and simulation of alternative treatment strategies to improve individual patient care, which would provide entirely new possibilities for clinical assessment and treatment adjustment. In addition, more fine-grained solutions to different exercise scenarios can be provided compared to current clinical guidelines that rely on observations of glucose changes during PA. We also anticipate that our model might find application in decision support systems or meal bolus calculators to determine insulin requirements for improved glycemic control, and would in particular allow to consider PA-induced changes in insulin sensitivity that can lead to late-onset hypoglycemia. Further applications might include the development of control algorithms for insulin treatment adjusted to glucose metabolism during and after exercise.

## Conclusion

We proposed a model of glucose-insulin regulation that captures the acute and prolonged effects of moderate-to high-intensity PA on glucose metabolism. The model accurately predicts BG during PA and subsequent recovery and is capable of describing data from individuals with T1D. We illustrated its use in replay simulations for personalized PA management in children, which could support clinicians in tailoring treatment strategies to individuals in the future. We also anticipate that it finds applications as an ‘exercise calculator’ [14] for clinical decision support, as well as for improving control algorithms for closed-loop insulin delivery.

We evaluated the model’s performance on several data sets, but further validation of the model and personalized replay are warranted before application in a clinical setting.

## Supporting information

**S1 File**

## Acknowledgments

The authors would like to thank Jörg Stelling for valuable feedback and discussions.

## Notes

### Competing Interest Statement

The authors have declared no competing interest.

